# Theoretical physical color gamuts define luminosity and naturalness perceptual limits in natural scenes

**DOI:** 10.1101/2025.04.04.647230

**Authors:** Killian Duay, Takehiro Nagai

## Abstract

Realism in augmented reality (AR) hinges on the seamless blending of virtual elements into real-world environments. One possible factor influencing this realism may be the physical gamut: an internal representation of all perceivable colors within a natural scene. Previous studies on luminosity thresholds suggest that this gamut, rooted in optimal colors theory, constrains perceptual judgments. While promising, such findings were based on abstract and two-dimensional stimuli only. Before extending this framework to more realistic AR scenarios, an essential next step is to assess whether the physical gamut theory also applies to naturalistic stimuli. This study addresses that gap. Our results reveal that the physical gamut remains a valid construct for natural objects viewed in realistic scenes. Moreover, observers’ judgments of luminosity thresholds appear guided not only by a criterion of self-luminosity, but also by an implicit sense of naturalness. These insights pave the way for exploring AR realism through the lens of physical gamut theory.

## Introduction

Augmented reality (AR) is defined by Billinghurst et al. as the seamless integration of virtual and real-world elements^1^. The ultimate goal of AR is to achieve the “total illusion”, where virtual objects are so realistic that users cannot discern them from reality. When this realism reaches a high level, individuals may experience a strong sense of presence in the virtual environment: a state in which the virtual world becomes more engaging than the physical surroundings, leading participants to perceive displayed scenes as places visited or real experiences lived, rather than as images seen^2^. Achieving this illusion involves numerous factors, which have been studied from both psychological and technological perspectives, including personality traits^3,4^, three-dimensional (3D) model precision^5^, texture quality^5,6^, or lighting and shadows fidelity^7–9^. While the role of lighting has primarily been investigated in terms of its effects on projected shadows, another key aspect influencing realism and presence could be how lighting affects the colors reflected by virtual objects and their perception by human observers, which has never been explored. A virtual object with lighting that does not match the real-world illumination may appear incongruous, either too bright for the scene or reflecting implausible colors, thereby breaking the realism and sense of presence. For instance, a pure white and highly luminous object in an orangish sunset scene may seem unnatural, as such a bright white would be physically impossible under these lighting conditions. This phenomenon aligns with the concept of a *physical gamut*, which encompasses all colors that can exist within a given context. A possible hypothesis is that humans have an internalized reference for this physical gamut, which they use to unconsciously assess the plausibility of objects or scenes, significantly influencing their sense of presence in both virtual and real environments. This raises the question of whether such a perceptual mechanism exists and plays a role in human visual processing when evaluating natural scenes, whether augmented or not.

Morimoto et al. demonstrated the potential existence of an internal representation of physical gamuts in the human brain in the context of distinguishing between self-luminous objects (e.g., light sources) and non self-luminous objects (e.g., surfaces reflecting light)^10^. The boundary between these categories is referred to as the luminosity thresholds. Their findings support the theory that when viewing an object in a scene, humans first estimate the light’s intensity and chromaticity, which they can do with moderate precision without being able to precisely identify its exact chromaticity or intensity^11–13^. Subsequently, the theory states that humans unconsciously determine whether the object’s color falls within or outside the internal representation of the physical gamut defined by the estimated light. If the object’s color lies outside the gamut, it surpasses the luminosity threshold and is perceived as self-luminous; otherwise, it remains within the gamut and appears as an illuminated surface. Their study explained that humans do not possess the physical gamut reference in a strictly physical sense within their brain, but rather construct an internal reference that approximates it. This internal representation is formed through empirical observations of the world and the surface colors encountered in daily life. Furthermore, their research revealed that the optimal colors form a highly accurate theoretical model for visualizing the internal representation of the physical gamut and closely matching it. Optimal colors are defined as surfaces with spectral reflectance values of either 0% or 100% at any given wavelength, exhibiting at most two abrupt spectral transitions^10,13,14^. In reality, physical constraints prevent any surface from achieving 100% reflectance at any wavelength, meaning that optimal colors inherently have higher luminance than any real surface with the same chromaticity. Therefore, no real surface can exceed this optimal colors distribution. By deriving all possible optimal reflectances and computing the spectral products with a given illuminant (i.e., light), the resultant set of optimal colors constitutes the physical gamut for that illuminant. Although optimal colors are not physically realizable, the authors demonstrated that they provide the most faithful model for visualizing the internal reference that humans seem to empirically tend to build. Overall, Morimoto et al. demonstrated a strong correlation and match between optimal colors and the luminosity thresholds, suggesting that the physical gamut theory accurately predicts these thresholds and that gamuts appear to be intrinsically grounded in optimal colors^10^. Specifically, optimal colors appear to shape and define the absolute theoretical boundaries of the physical gamut that humans tend to unconsciously reconstruct as internal representations based on empirical observation of the real world^10^. In other words, the physical gamut refers to the physical boundaries and constraints of colors in the real world and, although not directly analogous to the concept of a physical gamut, optimal colors currently provide the best model for computing and predicting these gamuts and the way humans internally represent them. For practical reasons, we will refer to this general concept as the “physical gamut” throughout the remainder of this study, although the more accurate term would be “an internal representation of the physical gamut”.

The concept of physical gamut could be applied to the problematic of presence in AR described in the first paragraph; if an object’s color falls outside the physical gamut in a given scene and the object is unlikely to be self-luminous (e.g., a cat or a flower), it will appear highly unrealistic in the given context, breaking the sense of presence. However, while this hypothesis is compelling, a critical gap remains in extending the physical gamut theory to AR scenarios: Morimoto et al. verified their model using abstract two-dimensional (2D) stimuli composed of flat and uniform dots^10^. These stimuli differ significantly from natural scenes and objects typically encountered in AR applications, making it uncertain whether their findings generalize to more complex natural environments. Indeed, prior research suggests that factors such as shading and texture significantly alter the perception of self-luminosity. For example, a uniform circle that appears self-luminous may seem half as self-luminous when presented as a shaded 3D sphere, and completely non self-luminous when additional surface texture is applied to the sphere^15^. More generally, shading is known to reduce perceived luminosity^16^. Furthermore, even minor surface details, such as a scratch, can eliminate the impression of self-luminosity and make an object appear as a standard reflective surface^15,16^. Therefore, prior to applying the physical gamut theory to the sense of presence in future innovative AR psychophysical experiments, it is necessary and appropriate to first verify whether the theory holds in more natural conditions than the abstract stimuli used by Morimoto et al.^10^ through a set of experiments in traditional vision science.

The present study aims to provide this intermediate missing piece by verifying whether the theory of physical gamut can also explain luminosity thresholds in more naturalistic stimuli. To this end, we conducted a series of three psychophysical experiments, drawing inspiration from the methodology of Morimoto et al.^10^, wherein observers adjusted the luminance of a target within a scene until it appeared self-luminous; these settings were then compared against predictions derived from the optimal colors model. The three experiments utilized natural scene background images but differed in the type of the target to adjust, increasing in naturalness across experiments: (1) a 2D uniform circle superimposed on the background in Experiment 1, (2) a shaded matte 3D sphere superimposed on the background in Experiment 2, and (3) an “in-picture” target in Experiment 3, where the pixels of an object within the original background image were directly manipulated. Based on aforementioned prior research, we hypothesized that the theory of the physical gamut would be: (1) confirmed in Experiment 1, as it has already been demonstrated that the background does not influence the results as long as it allows for an accurate estimation of the chromaticity and intensity of the illuminant^10^; (2) confirmed in Experiment 2, albeit with relatively higher observers’ settings than in Experiment 1 to compensate for the perceived reduction in luminosity induced by the shading of the sphere; and (3) refuted in Experiment 3, as the in-picture targets consist of natural objects with textures that disrupt the perception of self-luminosity. Remarkably and against all expectations, our results demonstrated that the physical gamut theory remained valid across all three experiments, successfully predicting and explaining luminosity thresholds in even the most naturalistic conditions. As anticipated, many observers reported that they could not directly perceive the targets as self-luminous in Experiments 2 and 3 but relied on an alternative strategy to assess luminosity thresholds: they judged them based on “unnatural brightness”. They reported that they were able to determine when the luminosity is excessively too high to be plausible, based on an impression of naturalness. Although they shifted their judgment criterion in order to be able to perform the task, their results were completely identical to those of observers who could directly judge self-luminosity. Moreover, the match with the physical gamut was observed equally well in both cases. These findings seem to indicate that the physical gamut theory is significantly more robust than previously assumed, with broader applications and ramifications extending beyond simply self-luminosity thresholds or 2D uniform circle luminosity judgments. The concept of physical gamut seems to apply to all types of images or scenes and definitively appears to be defined as “all possible colors in a natural scene under a given illuminant for non light-emitting objects”. It is intrinsically linked to the notion of luminosity thresholds which seem to encompass both self-luminosity thresholds and naturalness thresholds as similar and complementary concepts. These gamuts are encoded as internal references in the human visual system. These findings have profound potential implications for both applied fields (e.g., AR or projection mapping) and fundamental science (e.g., understanding human visual processing mechanisms). Having filled this critical gap, future research can now explore applying the physical gamut theory in AR to predict and evaluate virtual object realism and the participants’ sense of presence, potentially contributing to a deeper understanding of human visual perception in AR and supporting advancements in the technology.

We want to emphasize that, while we chose to build our study on top of the optimal colors theory and the concept of the physical gamut, other theories investigating luminosity thresholds could have been considered, including those based on: the highest intensity in the scene, the absolute intensity of the stimulus, the local or global contrast, or intensity comparisons with the average intensity of the scene^17^; a direct comparison with a white surface^18^; the purity of the stimulus or the chromaticity of the illumination^19–22^; or the brightness rather than the luminance of the stimulus^23^. Nonetheless, we chose to focus on the physical gamut, as this more recent theory has proven to be the most effective and superior to others, which fail to provide a thorough understanding of luminosity thresholds^10^. This does not imply that the theory is free from flaws or uncertainties, as previously noted; it is also the aim of the present study to examine it in greater depth.

## Results

### Experiment 1: the physical gamut theory holds for 2D uniform circle on top of natural backgrounds

In Experiment 1, eight observers were tasked with judging the luminosity thresholds of a target presented in different natural backgrounds. The target was a 2D uniform circle placed at the center of the background images (see Fig. 1c). Observers judged the luminosity thresholds for different chromaticities of the target across all natural backgrounds (see Fig. 1a and 1b). To determine the luminosity threshold, they were asked to adjust the luminance of the target using the scroll wheel of a mouse and set it to the point at which the target began to appear self-luminous (see Fig. 1e). The experiment was conducted under two different illuminants: once with stimuli generated under a neutral illuminant with a color temperature of 6500 Kelvin (K) and once under an orangish illuminant with a color temperature of 3000 K (see Fig. 1b and 1d). The aim of this experiment was to determine whether the observers’ settings matched the physical gamut defined by the optimal colors theory. To verify this, we plotted the average settings of all observers across all backgrounds for each chromaticity separately (forming the locus of observers’ settings) along with the optimal colors (forming the locus of optimal colors, or physical gamut). We then examined whether there was a correlation between the two loci. We plotted these data on a 2D diagram, selecting the CIE 1931 chromaticity coordinate *x* as the horizontal axis to improve readability of the charts. However, we want to emphasize that the concept of the physical gamut is inherently 3D, with the CIE 1931 chromaticity coordinates *x* and *y* forming the horizontal plane and luminance representing the elevation (see Fig. 1f). We hypothesized a strong correlation between the two loci, supporting the theory of physical gamut in these conditions. This expectation was based on prior research demonstrating that the theory holds for uniform circles against abstract backgrounds, provided that the backgrounds allow for the identification of the chromaticity and intensity of the illuminant^10^. Given that natural backgrounds contain rich illumination cues, they were expected to support this effect.

**Figure 1.**
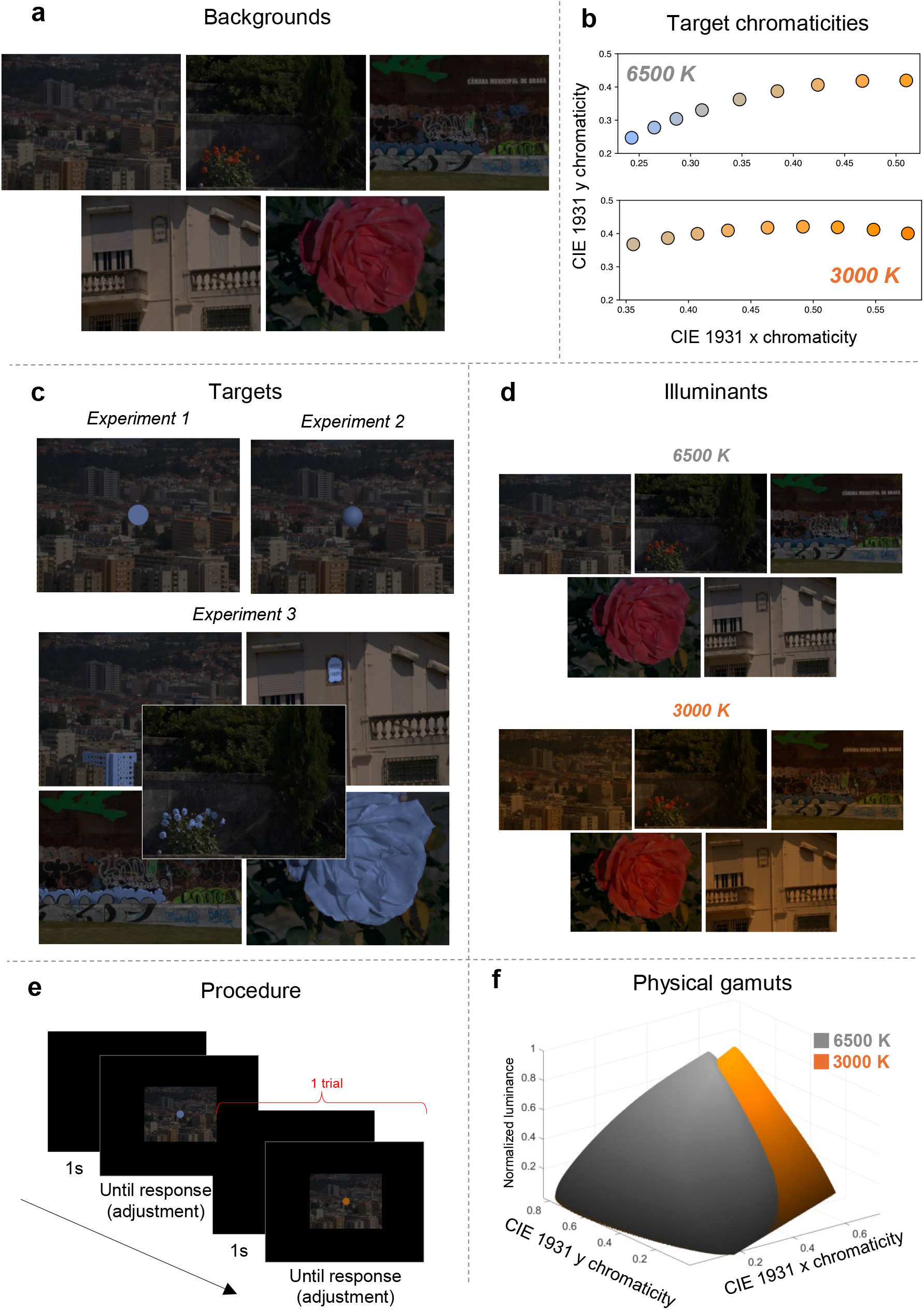
Methods of the three experiments. **a**: backgrounds of the experiments. **b**: test chromaticities of the targets. **c**: targets of Experiments 1,2, and 3. **d**: illuminants of the experiments. **e**: procedure of the experiments with an example of two trials. **f**: normalized physical gamuts of illuminants 6500 K and 3000 K.

As anticipated, the results confirmed the validity of the physical gamut theory for 2D uniform circles on natural backgrounds. Figure 2a shows the observers’ settings and optimal colors loci separately for the 6500 K and 3000 K illuminant conditions. A strong correlation was found between the two loci (for 6500 K: *r* = 0.87, *p* < 0.001; for 3000 K: *r* = 0.76, *p* < 0.001). Additionally, the shapes of the loci are similar; in other words, the perceived luminosity thresholds match the theoretical physical gamut derived from optimal colors. Consistent with Morimoto et al.^10^, the chromaticity of the observers’ peak setting corresponds to the chromaticity of the white point of the illuminant. Indeed, a shift in peak settings was observed between the 6500 K and 3000 K conditions, following the white point of the illuminant. Furthermore, as reported by Morimoto et al.^10^, the optimal colors theory appears to be slightly less precise for the illuminant 3000 K. This discrepancy can be attributed to the theory’s reliance on the estimation of the chromaticity and intensity of the illuminant. Since estimating an orangish illuminant is inherently more challenging and less precise than estimating a neutral illuminant (which corresponds to 6500 K in our case), the reduced accuracy for 3000 K can be expected^10,12,13^.

**Figure 2.**
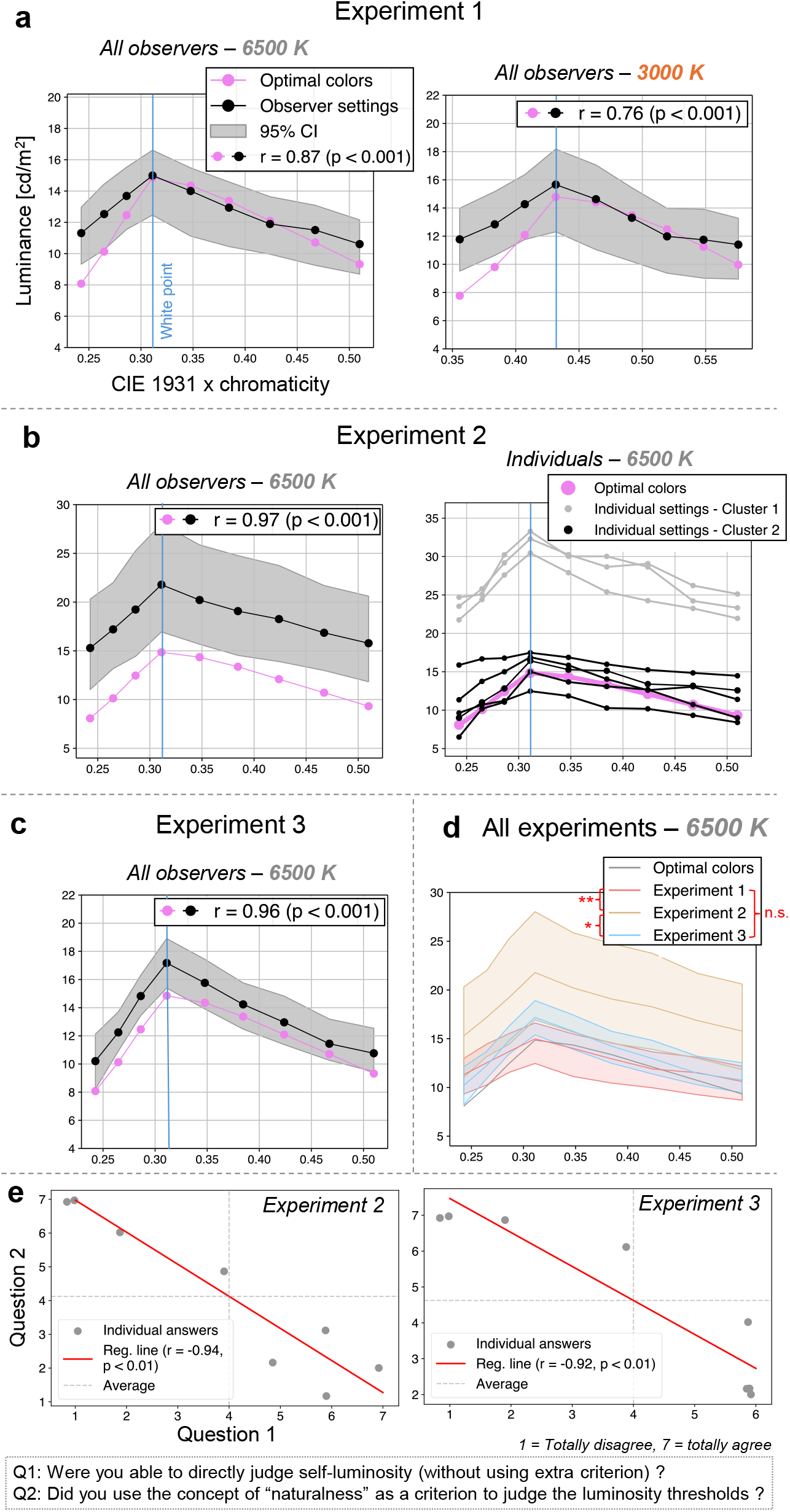
Results of the experiments. **a**: results of Experiment 1 (averaged across all observers and backgrounds for condition 6500 K on the left and 3000 K on the right). **b**: results of Experiment 2 (averaged across all observers and backgrounds on the left and individual results on the right). **c**: results of Experiment 3 averaged across all observers and backgrounds. **d**: results of the three experiments averaged across all observers and backgrounds plotted on the same chart (only condition with illuminant 6500 K for Experiment 1). **e**: results of the second questionnaire for Experiment 2 (on the left) and Experiment 3 (on the right). Asterisks indicate significant differences between the loci: * *p* <0.05 and ** *p* <0.01. Filled colored areas represent 95% confidence intervals. See the Methods section for details on the statistical analysis.

### Experiment 2: the physical gamut theory holds for 3D sphere on top of natural backgrounds

The design, task, and objective of Experiment 2 were similar to those of Experiment 1, with the primary difference being that the target was a 3D sphere instead of a 2D uniform circle. The aim here was to enhance the naturalness of the stimulus compared to Experiment 1 and to assess whether the physical gamut theory still holds. To create the impression of three-dimensionality, shading was applied to the target from Experiment 1: while the maximum luminance was preserved for a portion of the circle, a circular and directional gradient was introduced to the rest of the surface (see Fig. 1c). Two additional modifications were introduced in Experiment 2: only the neutral illuminant was used and a questionnaire was administered at the end of the experiment to evaluate the observers’ subjective experience regarding task difficulty and collect spontaneous comments. The questions and details of the questionnaire are presented in Supplementary Table S1 (see Supplementary Information). We hypothesized that the theory of physical gamuts would be validated in Experiment 2, as in Experiment 1. However, based on previously introduced studies, we anticipated that observers’ settings would be relatively higher than in Experiment 1 to compensate for the perceived reduction in luminosity induced by the shading applied to the 3D sphere.

As expected, the theory of physical gamuts was confirmed for 3D spheres presented against natural backgrounds. Figure 2b shows the loci of the observers’ settings and the optimal colors, plotted both as an average across all participants and as individual data points. As in Experiment 1, a strong correlation was observed between the optimal colors loci and the averaged observers’ settings (*r* = 0.97, *p* < 0.001), with both exhibiting similar shapes. Moreover, the peak of the observers’ settings corresponds to the white point of the illuminant. However, the magnitude of the settings is higher in Experiment 2 than in Experiment 1, with an average peak setting of 21.8 compared to 15.0 in Experiment 1. Additionally, the confidence interval is broader in Experiment 2 (range of 11.1) than in Experiment 1 (range of 4.15) for the highest setting, which corresponds to the white point. We assessed intra-observer variability in Experiment 2 and obtained an overall confidence interval with a range of 5.67 (for each observer, confidence intervals were calculated individually for each chromaticity, then averaged to yield a per-observer mean, and finally averaged across all observers to obtain the overall interval). These findings suggest greater inter-observer variability when assessing the 3D sphere. Upon closer examination of individual results (see Fig. 2b), two distinct observer groups appeared to emerge. To confirm this, we performed a clustering analysis: the optimal number of clusters was determined using the Elbow method, followed by k-means clustering. This analysis confirmed the existence of two well-separated observer groups, as indicated by the Silhouette Score (0.79) and the Davies-Bouldin Index (0.24). Our interpretation of these clusters suggests that observers in cluster 2 judged the luminosity thresholds based on the highest luminance of the sphere. Their peak setting averages 15.6, closely aligning with the average peak setting from Experiment 1 (15.0). Given that the maximum luminance of the 3D sphere matched that of the 2D uniform circle in Experiment 1, it appears that these observers based their judgments on the brightest portion of the sphere. In contrast, observers in cluster 1 seemed to assess the luminosity thresholds based on the sphere’s average luminance. Their average peak setting is 32, approximately twice those of cluster 2 or Experiment 1. This suggests that these observers compensated for the reduced perceived luminosity caused by shading. Indeed, due to the shading applied to simulate three-dimensionality, the average luminance of the sphere was approximately 50% of the original luminance of the 2D uniform circle in Experiment 1. Thus, these observers appeared to base their judgments on average luminance and adjusted their settings to compensate for the 50% reduction by doubling their original settings. The questionnaire results (see Supplementary Table S1 in Supplementary Information) support this interpretation. The eight observers in the experiment responded to this questionnaire. Overall, the task was reported as difficult. Three observers specifically mentioned a “gray zone” between the “illuminated surface” and “self-luminous” appearance, and they noted that this gray zone was significantly wider than in Experiment 1. This supports the idea that different perceptual strategies were employed: the ambiguity of this gray zone may reflect whether observers based their judgments on the sphere’s highest luminance or its average luminance. Observers ultimately made a choice in how they interpreted this perceptual boundary. Additionally, five observers reported that the sphere never appeared self-luminous and three observers reported that they were still able to judge the self-luminosity thresholds based on perceptual naturalness rather than strict self-luminosity. At high luminance levels, the sphere appeared unnatural, as it was perceived as being relatively too bright to be physically plausible in the given scene; this was the breaking point they used for their judgment. Interestingly, although these observers did not perceive the stimuli as self-luminous and instead judged them based on “unnatural brightness”, their results were comparable to those of other participants who did perceive self-luminosity.

### Experiment 3: the physical gamut theory holds for in-picture objects in natural backgrounds

The design, task, and objective of Experiment 3 were similar to those of Experiments 1 and 2. However, the key difference was that the target in this experiment was an “in-picture” element, meaning it was an object already present within the original background rather than a superimposed stimulus (such as the uniform circle in Experiment 1 or the 3D sphere in Experiment 2; see Fig. 1c where the targets of Experiment 3 are highlighted in light blue). The aim here was to further enhance the naturalness of the stimulus compared to Experiments 1 and 2 and to evaluate whether the physical gamut theory still holds. In addition, as in Experiment 2, only the neutral illuminant was used, and a questionnaire was administered at the end of the experiment to assess observers’ subjective experience regarding the task’s difficulty and collect spontaneous comments. The questions and details of the questionnaire are presented in Supplementary Table S2 (see Supplementary Information). Based on previously introduced studies, our hypothesis was that the theory of physical gamuts would not hold in Experiment 3. We anticipated that, because the in-picture targets were natural objects with textures, they would disrupt the perception of self-luminosity, preventing the same results observed in Experiments 1 and 2 from emerging.

Contrary to expectations, the results revealed that the theory of physical gamuts was fully validated even for in-picture targets embedded within natural backgrounds. Figure 2c illustrates the loci of the observers’ settings and the optimal colors, plotted as an average across all participants. As in Experiments 1 and 2, a strong correlation was observed between the optimal colors loci and the observers’ settings (*r* = 0.96, *p* < 0.001), with both exhibiting a similar shape. Furthermore, the peak of the observers’ settings once again corresponded to the white point of the illuminant. However, in contrast to Experiment 2, the magnitude of the settings in Experiment 3 was not notably higher than in Experiment 1 (average peak setting of 17.1 in Experiment 3 vs. 15.0 in Experiment 1). Likewise, the confidence interval was not notably broader (range of 3.57 in Experiment 3 vs. 4.15 in Experiment 1). This suggests that, unlike in Experiment 2, observers did not adopt different strategies for judging luminosity thresholds (i.e., judging based on the average luminance of the target vs. its maximum luminance). A plausible explanation is that, in Experiment 3, the target was an integral part of the original image, allowing observers to make direct comparisons with the surrounding elements in the scene. The questionnaire results (see Supplementary Table S2 in Supplementary Information) further support this interpretation. The eight observers in the experiment responded to this questionnaire. The task was reported to be markedly easier than in Experiment 2. Four observers explicitly stated that the task was straightforward because the target was part of the original scene, enabling them to compare it directly with the background. Moreover, as in Experiment 2, four observers reported that the target never appeared self-luminous, and three observers reported that they were still able to judge the self-luminosity threshold based on perceptual naturalness. They reported that, at high luminance levels, the target appeared unnatural, as it was perceived as being relatively too bright to be physically plausible in the given scene; this was the breaking point they used for their judgment. Again, although these observers did not perceive the stimuli as self-luminous and instead judged them based on “unnatural brightness”, their results were comparable to those of other participants who did perceive self-luminosity. In addition, the same observers reported that this unnaturalness was even easier to detect. Indeed, when the target’s luminance increased beyond a certain threshold, it appeared to detach from the original image, creating the impression that it was no longer part of its original plane but instead shifted closer to the observer’s eyes. This effect facilitated the identification of the point at which the target became “too bright” or “unnatural”. This phenomenon might also explain why, unlike in Experiment 2, observers did not split into two groups based on different perceptual strategies (i.e., judging based on the average luminance of the target vs. its maximum luminance). Because the target was part of the original image, observers could directly compare it with the background, making judgments about unnaturalness or self-luminosity more straightforward and consistent across participants.

## Discussion

For Experiment 1, our hypothesis was that the physical gamut theory would hold in natural backgrounds with targets in the form of 2D uniform circles. This hypothesis was confirmed for two different illuminants, with the theory performing slightly better under neutral light compared to orangish light, reinforcing previous findings under slightly different conditions. For Experiment 2, we hypothesized that the physical gamut theory would be valid for natural backgrounds with 3D spherical targets, but that observers would set higher thresholds to compensate for shading effects on the sphere. This was only partially confirmed: some observers indeed set thresholds twice as high to compensate for shading over 50% of the sphere’s surface, while others made settings identical to those in Experiment 1. For Experiment 3, we hypothesized that the physical gamut theory would not hold, as textured natural objects are never perceived as self-luminous. However, contrary to expectations, this hypothesis was entirely refuted, as the physical gamut theory was fully validated in this scenario, if not more robustly than in Experiment 1. Figure 2d summarizes these results and confirms that the locus of observers’ settings was significantly higher in Experiment 2 than in Experiment 1 (*p* < 0.01) and Experiment 3 (*p* < 0.05). The loci of observers’ settings in Experiment 1 and Experiment 3 did not differ significantly (*p* = 0.37).

These results, in conjunction with the questionnaire data, indicate that the physical gamut theory is more robust than previously anticipated and extends the concept of the physical gamut beyond merely predicting self-luminosity thresholds. Many observers reported that they could not perceive the targets as self-luminous; instead, they judged them based on a concept of “unnatural brightness”, yet their results were comparable to those of participants who did perceive self-luminosity. Therefore, when observers are asked to assess the luminosity thresholds of objects that they do not perceive as self-luminous, they appear to rely on the notion of unnaturalness (defined, based on observers’ comments, as “a luminosity level too high for the scene to be plausible”). The physical gamut theory appears to encompass and define these unnaturalness thresholds alongside self-luminosity thresholds. This suggests that self-luminosity and unnaturalness thresholds share a common conceptual framework, defining a general physical gamut that extends its definition to “all possible colors in a scene for an object that does not emit light”. In other words, it seems that luminosity thresholds are not strictly self-luminosity thresholds but instead incorporate both self-luminosity and unnaturalness thresholds, which are conceptually distinct yet can be interpreted similarly. After analyzing the responses from the first questionnaire associated with Experiments 2 and 3, we conducted a second questionnaire to verify this conclusion. The goal of the follow-up questionnaire was to determine whether all observers were unable to directly assess self-luminosity and whether they instead defaulted to a concept of unnaturalness. The details of the questions and results are presented in Fig. 2e. The results indicate a strong negative correlation between the ability to directly assess the self-luminosity of targets and the reliance on naturalness as a judgment criterion (for Experiment 2: *r* = -0.94, *p* < 0.01; for Experiment 3: *r* = -0.92, *p* < 0.01). When observers can directly evaluate self-luminosity, they do not rely on the concept of naturalness.

However, when they are unable to make this direct assessment, they use naturalness as a reference. The findings suggest that approximately one-third of observers are clearly able to judge self-luminosity and make little or no use of naturalness, another third struggle to assess self-luminosity and rely predominantly on naturalness, while the remaining third fall in between, integrating both aspects into their judgments. Nevertheless, interestingly, the settings aligned with the physical gamut defined by optimal colors for all observers, regardless of how they judged the thresholds. This seems to confirm that the luminosity thresholds shaping the physical gamut, derived from optimal colors, apply equally to self-luminosity and unnaturalness concepts. These two concepts appear to be intrinsically linked, differing in semantics (i.e., “an object being a light source” vs. “an object appearing too bright for the scene”) but not in their absolute threshold values, which remain identical, or interpretation, which are similar.

Furthermore, whether observers directly judged self-luminosity or relied on unnaturalness, they appeared to use two distinct strategies to assess the luminosity thresholds of more complex natural stimuli (3D spheres): either by considering the target’s average luminance or its maximum luminance. This introduced difficulty, as observers needed to arbitrate between these two strategies, creating ambiguity. However, this difficulty was absent when complex natural objects were directly embedded into the original scene (in-picture targets), as observers could compare them directly to the background. Many observers reported that targets appeared to “come out” of the image when their luminance was too high relative to their surroundings, reinforcing a sense of unnaturalness or self-luminosity and making the task easier by eliminating the need for strategy arbitration. This direct comparison to the local surroundings and the effect of the target coming out of the image was not present in Experiment 2, where the 3D sphere, being superimposed onto the background via computer graphics, was already perceived as separate from the image. This distinction might explain why two different strategies were used in Experiment 2 but not in Experiment 3. This observation also raises important questions in other experimental contexts, such as color constancy. For instance, can stimuli involving 3D spheres or 2D uniform circles superimposed on natural backgrounds be effectively used to evaluate color constancy if they are perceived as separate from the original scene and illumination ?

Finally, one might note that the results from Experiment 3 were superior to those from Experiments 1 and 2 in terms of both the correlation between loci and the magnitude of the distance between them. Based on our previous conclusions, we attribute this to the fully natural composition of Experiment 3, which made judgments significantly easier. Whether assessing self-luminosity directly or relying on unnaturalness, observers could use the background and local surroundings for direct comparison. In contrast, in Experiments 1 and 2, targets superimposed via computer graphics introduced ambiguity and increased task difficulty, as the target was not necessarily perceived as an integral part of the background. Similarly, Morimoto et al. obtained results superior to those of our Experiment 1, but their stimuli consisted entirely of 2D uniform circle patterns (both background and target), allowing targets to be perceived as embedded within the background due to their uniform depth and planar alignment^10^. This supports the notion that mixed stimuli, combining real backgrounds with computer-generated targets, introduce ambiguity in judgments; a factor absent in fully natural or fully abstract stimuli, leading to a task easier to conduct in these conditions and superior results. These results, along with the observations described in the previous paragraph, suggest that the consistency and naturalness of an image modulate the way a stimulus is perceived and may influence task performance in psychophysical experiments.

Our study has several limitations that future research could address. First, due to time constraints, we used a limited set of backgrounds, illuminations, and target chromaticities. Morimoto et al. demonstrated that the physical gamuts are less accurately predictive when target chromaticities or illuminations are highly unnatural, such as green or magenta^10^. Second, due to the already extensive duration of the experiments, we tested only one illumination color in Experiments 2 and 3. This choice was guided by findings from both Morimoto et al.^10^ and our own Experiment 1, which already support extensively the validity of the physical gamut theory under different illumination conditions and suggest that observers adapt the gamuts to the illumination color. We considered unnecessary to re-demonstrate it in Experiments 2 and 3. Despite these first two limitations, the focus of the present study was on evaluating the physical gamut theory using more naturalistic stimuli; we determined that the combinations of target chromaticities and illuminations were sufficient to test our hypotheses. Nonetheless, future studies could verify whether the physical gamut theory holds across a greater diversity of backgrounds, target chromaticities, and illuminant colors, as this remains unclear. Third, we have focused solely on the optimal colors theory and the concept of the physical gamut to explain and define the luminosity thresholds. We took this stance because it is the most recent theory, and it has been shown to be the most effective and superior to other theories^10^. However, future studies could explore other previous theories that explained luminosity thresholds based on: the highest intensity in the scene, the absolute intensity of the stimulus, the local or global contrast, or intensity comparisons with the average intensity of the scene^17^; a direct comparison with a white surface^18^; the purity of the stimulus or the chromaticity of the illumination^19–22^; or the brightness rather than the luminance of the stimulus^23^. Finally, we tested our theory with only eight observers. It would be valuable to assess these findings in a larger sample, particularly to further validate the different judgment strategies proposed in this study (i.e., average vs. maximum luminance of the target and self-luminosity vs. unnaturalness).

In conclusion, we have demonstrated that the physical gamut theory is significantly more robust than previously anticipated, with broader applications beyond the strict definition of self-luminosity thresholds and 2D uniform circle judgments. Through different judgment strategies (i.e., average vs. maximum luminance of the target and self-luminosity vs. unnaturalness), the concept of the physical gamut appears to apply universally to all types of stimuli and a representation of it may be fundamentally encoded in the human brain. The notion of physical gamut definitely seems to approach a definition such as “all possible colors in a scene for an object that is not a light”. The self-luminosity and unnaturalness thresholds shaped by the physical gamut seem intrinsically linked, resting on the same luminosity threshold principles. These findings have potential implications for both applied fields (e.g., AR, projection mapping) and fundamental science (e.g., understanding human visual mechanisms). Now that we have filled this gap and shown that the physical gamut hold for natural stimuli through traditional vision science experiments, future studies could explore applying these principles in AR to predict and assess the realism and presence of virtual objects in innovative psychophysical experiments. Indeed, we believe that the physical gamut concept governing impressions of object naturalness in 3D scenes, as revealed in this study, could provide significant insights for evaluating realism and presence in AR and significantly contribute to the advancement of the technology.

## Methods

### Observers

The observers were two females and six males. All were undergraduate or graduate students from Institute of Science Tokyo. All observers passed the Ishihara color vision test and had normal or corrected-to-normal visual acuity. All experiments were designed in accordance with the Declaration of Helsinki and was approved by the Ethical Review Committee of Institute of Science Tokyo. Informed written consent was obtained from all observers after explaining the details of experimental protocols.

### Apparatus

The experiments took place in a darkroom. Stimuli were displayed on an OLED monitor (PVM-A250, Sony, Japan) with a 10-bit per channel color resolution, a spatial resolution of 1920 × 1080 pixels, a 25-inch screen diagonal, and a 60 Hz refresh rate. To ensure accurate luminance and chromaticity presentation, the display was carefully calibrated using a spectroradiometer (Specbos1211-2, JETI Technische Instrumente GmbH, Germany) and a colorimeter (ColorCAL II, Cambridge Research Systems, UK). The monitor was connected to a laptop (MacBook Air, Apple, USA; Apple M2, macOS Sequoia 15.2), and the experiments were controlled by a custom program written in PsychoPy 2024.2.4. Observers’ heads were stabilized using a chinrest at a 40 cm viewing distance, and responses were collected via a numeric keypad and a mouse.

### Color computation

In color science, the CIE 1931 color space provides a standardized method to quantify color perception based on human vision. To compute colors in this space, three key components are required^24^: the spectral reflectance of a surface, the spectral power distribution (SPD) of the illuminant, and the CIE 1931 color matching functions (CMF), which represent the average human eye’s sensitivity to different wavelengths (*2° Standard Observer* in the present study). The spectral reflectance describes how much light a surface reflects at each wavelength, while the illuminant’s SPD defines the intensity of incident light across the spectrum. The reflected light is computed by multiplying the reflectance with the illuminant at each wavelength. This product is then integrated across the visible spectrum, weighted by the CMFs, to yield the final color expressed in tristimulus values *(X, Y, Z)*, from which the chromaticity coordinates *(x*,*y)* and the luminance *Y* can be derived. Hyperspectral images contain detailed spectral reflectance information for each pixel, enabling accurate per-pixel color computation under any defined illuminant. For more details, see previous literature^24,25^. This is the method we used to generate the colors and stimuli in this study.

### Stimulus

The stimulus was presented at the center of a completely black background of 0 candelas per square meter (cd/m^2^), made possible by the OLED display used in our setup, as illustrated in Fig. 1e. The stimulus subtended approximately 13.26° horizontally and 10.00° vertically, and was composed of two main elements: a background and a target.

Five different backgrounds were used (see Fig. 1a), each derived from hyperspectral images obtained from a publicly available dataset^26^. These images were selected to span a variety of natural scenes, including urban environments, vegetation-rich areas, and mixed-content settings, thereby sampling diverse distributions of natural colors. To simulate different lighting conditions, we applied two illuminants corresponding to distinct correlated color temperatures (CCTs): a neutral daylight-like illuminant (6500 K) and a warmer, orangish illuminant (3000 K). The backgrounds were generated by multiplying the hyperspectral data with the spectral power distributions of Planckian radiators at 6500 K and 3000 K, respectively, as shown in Fig. 1d. The illuminant intensities were scaled such that a perfect white surface (reflecting 100% of all wavelengths) under either lighting condition would yield a luminance of 15 cd/m^2^. Importantly, the same set of backgrounds was used across the three experiments.

The targets differed between the three experiments, as shown in Fig. 1c. In Experiment 1, the target was a 2D uniform circle with a diameter of approximately 1.29°, overlaid on the background. In Experiment 2, the target was a 3D sphere subtending the same visual angle. To simulate three-dimensionality, a shading gradient was applied to the 2D circle from Experiment 1: while maintaining the maximum luminance on one portion of the surface, a circular and directional luminance gradient was introduced over the remainder (see Supplementary Fig. S1 in Supplementary Information). In Experiment 3, the targets were real-world objects originally present within the backgrounds. These objects, highlighted in blue in Fig. 1c, were isolated and manipulated using masks. They were chosen to represent a wide range of physical properties, including differences in semantic category (e.g., buildings or flowers) and spatial extent (occupying either large or small portions of the image).

The target colors were defined by a chromaticity and luminance values in CIE 1931 color space. Across all conditions, the target could assume one of nine distinct chromaticities per trial, as illustrated in Fig. 1b. Only one chromaticity was applied at a time, and it was uniformly mapped onto the target (as exemplified by the blue chromaticity in Fig. 1c; the chromaticity coordinates *(x,y)* of these pixels were transformed into the test chromaticities). Following a strategy similar to that of Morimoto et al.^10^ to adopt natural colors, the chromaticities were selected to lie along the black body locus and corresponded to Planckian radiator CCTs of 2000 K, 2400 K, 2900 K, 3500 K, 4300 K, 5600 K, 7200 K, 10000 K, and 20000 K. These were chosen to be approximately equally spaced along the locus on the CIE 1931 chromaticity diagram. The spectral power distributions of these nine Planckian radiators were multiplied with the spectral distributions of the two illuminants used in the experiments (3000 K and 6500 K), yielding the final test chromaticities. Notably, the chromaticity resulting from the 5600 K Planckian radiator closely matched the white point of both experimental illuminants.

The target luminance was directly set and adjusted by the observers’ during each trial task. The method for applying the observer’s selected luminance value to the target varied across the three experiments. In Experiment 1, the luminance was applied uniformly to all pixels of the 2D circular target. In Experiment 2, the selected luminance was applied to each pixel of the sphere and multiplied by the shading, such that the area of maximum luminance on the sphere corresponds to the value chosen by the observer, then gradually decreases according to the shading. In Experiment 3, the luminance adjustment was applied to the masked region corresponding to the target object. To preserve the object’s internal structure, the adjusted luminance was multiplied by the original pixel luminance values, which had been normalized between 0 and 1 such that the highest pixel value matched the observer’s chosen luminance.

All resulting stimuli were generated in the CIE 1931 color space and then converted to corresponding RGB values for the display used in our experiment. Final color rendering on the monitor was ensured through a color calibration procedure, based on prior spectral measurements and display characterization.

### Procedure

Figure 1e illustrates two example trials from the experimental procedure. Each trial began with a 1-second blank and black screen, followed by the presentation of the stimulus, which remained visible until the observer judged their task to be complete and validated their response using the numeric keypad. The task consisted of adjusting the luminance of the target using a combination of mouse controls. Specifically, the mouse scroll wheel allowed for luminance changes in steps of 1.0 cd/m^2^, while the left and right mouse buttons provided finer adjustments, decreasing or increasing luminance in 0.5 cd/m^2^ increments, respectively. The observer’s goal was to determine and set the luminance at which the target began to appear self-luminous.

During a trial, only the luminance of the target was modified; the chromaticity remained fixed. Technically, this was achieved by converting the RGB representation of the stimulus back into the CIE 1931 color space, adjusting the luminance component, and converting it back into RGB for display. This process was performed in real-time and was imperceptible to the observer, ensuring a seamless visual experience.

Each session consisted of 45 trials, corresponding to all combinations of five background scenes and nine target chromaticities. The order of the five backgrounds was randomized, and within each background block, the chromaticities were presented in random order. Importantly, each session was associated with a single illuminant condition and did not mix stimuli from different illuminant sets. For example, a session dedicated to the illuminant 6500 K featured only the five backgrounds and nine chromaticities computed under that specific illuminant.

In Experiment 1, observers completed four sessions under the 6500 K illuminant and four under the 3000 K illuminant, in randomized order. In Experiments 2 and 3, only the 6500 K illuminant was used, and observers completed four sessions for each experiment. Observers were permitted to take a break between sessions.

Each session began with a 2-minute dark adaptation phase during which the screen remained entirely black, followed by a 30-second adaptation phase to the experimental illuminant and background content. During this adaptation period, the five background images used in the current session were displayed in a continuous loop, each shown for two seconds.

Experiment 1 was conducted on a single day. Experiments 2 and 3 were performed on a different day, with a 30-minute break between them. At the conclusion of Experiments 2 and 3, observers were asked to complete a questionnaire regarding their perceptual experience and feelings, as detailed in Supplementary Tables S1 and S2 (see Supplementary Information).

### Physical gamut and optimal colors

Optimal colors are defined as surfaces whose spectral reflectance functions consist exclusively of 0% and 100% reflectance values across the wavelength spectrum, with at most two abrupt transitions between these extremes^10,13,14^. In the real world, however, no material can achieve 100% reflectance at any wavelength due to physical limitations. Consequently, optimal colors exhibit higher luminance than any real surface possessing the same chromaticity, and thus, no naturally occurring surface can exceed the luminance of these theoretical distributions at a given chromaticity. Therefore, by computing all possible optimal reflectance functions and multiplying them spectrally with a given illuminant, one obtains the complete set of optimal colors that form the physical gamut for that illuminant. In our study, we generated optimal colors in the CIE 1931 color space following established methodologies^10,13,14^. Specifically, we produced approximately 90,000 optimal spectral reflectance functions, corresponding to every possible combination of reflectance transitions that satisfy the criteria of optimal colors. Each spectrum was then multiplied with the spectral power distributions of the Planckian radiators at 6500 K and 3000 K. The resulting sets of optimal colors, forming the physical gamuts under each illuminant condition, are shown in Fig. 1f. The specific optimal colors corresponding to the chromaticities of the experimental targets, used for plotting and analyzing the 2D loci in Fig. 2, were selected from within this set. It is important to note that the physical gamut displayed in Fig. 1f is expressed in normalized luminance values. To obtain absolute luminance, these values must be multiplied by the scene’s illuminant intensity, which in our experiments was fixed at 15 cd/m^2^.

### Analysis

All statistical tests, confidence intervals, and *p*-values reported in this study were performed using a two-tailed, non-parametric bias-corrected and accelerated (BCa) bootstrapping method with 10,000 resamples of the eight observers and a significance level of 5%^27^. This approach provides robust estimates that account for potential bias and skewness in the sampling distribution. All *r*-values reported, including those associated with the regression lines in Fig. 2e, correspond to Pearson’s correlation coefficients. The statistical significance of these correlations (*p*-values) was assessed using the aforementioned BCa bootstrapping procedure. The regression lines presented in Fig. 2e were obtained via standard linear least-squares regression. The accompanying *r*-values represent the Pearson’s correlation coefficients and their significance levels were likewise computed using the aforementioned BCa method.

## Data availability

All data generated or analysed during the current study are available from the corresponding author on reasonable request.

## Additional Information: Competing interests

The authors declare no competing interests.

## Acknowledgements

This study was supported by JST SPRING, Grant Number JPMJSP2180 to KD, and JSPS KAKENHI Grant Number 23K28174 and 21KK0203 to TN.

## Author contributions statement

KD and TN designed the experiment, interpreted the results, and wrote the manuscript. KD conducted the experiments and analyzed the results. All authors reviewed and approved the final version of the manuscript.

## References

1. Billinghurst, M., Clark, A., Lee, G. et al. A survey of augmented reality. Foundations Trends Human–Computer Interact. 8, 73–272 (2015).

2. Slater, M. & Wilbur, S. A framework for immersive virtual environments (five): Speculations on the role of presence in virtual environments. Presence: Teleoperators & Virtual Environ. 6, 603–616 (1997).

3. Wallach, H. S., Safir, M. P. & Samana, R. Personality variables and presence. Virtual Real. 14, 3–13 (2010).

4. Nicovich, S. G., Boller, G. W. & Cornwell, T. B. Experienced presence within computer-mediated communications: Initial explorations on the effects of gender with respect to empathy and immersion. J. Comput. Commun. 10, JCMC1023 (2005).

5. Hvass, J. et al. Visual realism and presence in a virtual reality game. In 2017 3DTV conference: The true vision-capture, Transmission and Display of 3D video (3DTV-CON), 1–4 (IEEE, 2017).

6. Zimmons, P. & Panter, A. The influence of rendering quality on presence and task performance in a virtual environment. In IEEE Virtual Reality, 2003. Proceedings., 293–294 (IEEE, 2003).

7. Kán, P., Dünser, A., Billinghurst, M., Schönauer, C. & Kaufmann, H. The effects of direct and global illumination on presence in augmented reality. In Challenging Presence-Proceedings of 15th International Conference on Presence (ISPR 2014), 223–230 (Facultas Verlags-und Buchhandels AG, 2014).

8. Sugano, N., Kato, H. & Tachibana, K. The effects of shadow representation of virtual objects in augmented reality. In The Second IEEE and ACM International Symposium on Mixed and Augmented Reality, 2003. Proceedings., 76–83 (IEEE, 2003).

9. Tuceryan, M. et al. Cubemap360: interactive global illumination for augmented reality in dynamic environment. In 2019 SoutheastCon, 1–8 (IEEE, 2019).

10. Morimoto, T., Numata, A., Fukuda, K. & Uchikawa, K. Luminosity thresholds of colored surfaces are determined by their upper-limit luminances empirically internalized in the visual system. J. vision 21, 3–3 (2021).

11. Fukuda, K. & Uchikawa, K. Color constancy in a scene with bright colors that do not have a fully natural surface appearance. J. Opt. Soc. Am. A 31, A239–A246 (2014).

12. Morimoto, T., Fukuda, K. & Uchikawa, K. Effects of surrounding stimulus properties on color constancy based on luminance balance. J. Opt. Soc. Am. A 33, A214–A227 (2016).

13. Morimoto, T., Kusuyama, T., Fukuda, K. & Uchikawa, K. Human color constancy based on the geometry of color distributions. J. Vis. 21, 7–7 (2021).

14. Uchikawa, K., Fukuda, K., Kitazawa, Y. & MacLeod, D. I. Estimating illuminant color based on luminance balance of surfaces. J. Opt. Soc. Am. A 29, A133–A143 (2012).

15. Kuriki, I. Effect of material perception on mode of color appearance. J. vision 15, 4–4 (2015).

16. Kuriki, I. & Uchikawa, K. Limitations of surface-color and apparent-color constancy. J. Opt. Soc. Am. A 13, 1622–1636 (1996).

17. UlIman, S. On visual detection of light sources. Biol. cybernetics 21, 205–212 (1976).

18. Bonato, F. & Gilchrist, A. L. The perception of luminosity on different backgrounds and in different illuminations. Perception 23, 991–1006 (1994).

19. Evans, R. M. Fluorescence and gray content of surface colors. J. Opt. Soc. Am. 49, 1049–1059 (1959).

20. Evans, R. M. & Swenholt, B. K. Chromatic strength of colors: dominant wavelength and purity. J. Opt. Soc. Am. 57, 1319–1324 (1967).

21. Evans, R. M. & Swenholt, B. K. Chromatic strengths of colors, part ii. the munsell system. J. Opt. Soc. Am. 58, 580–584 (1968).

22. Evans, R. M. & Swenholt, B. K. Chromatic strength of colors, iii. chromatic surrounds and discussion. J. Opt. Soc. Am. 59, 628–634 (1969).

23. Uchikawa, K., Koida, K., Meguro, T., Yamauchi, Y. & Kuriki, I. Brightness, not luminance, determines transition from the surface-color to the aperture-color mode for colored lights. J. Opt. Soc. Am. A 18, 737–746 (2001).

24. Wyszecki, G. & Stiles, W. S. Color science: concepts and methods, quantitative data and formulae (John wiley & sons, 2000).

25. Foster, D. H. & Reeves, A. Colour constancy failures expected in colourful environments. Proc. Royal Soc. B 289, 20212483 (2022).

26. Foster, D., Amano, K. & Nascimento, S. Fifty hyperspectral reflectance images of outdoor scenes, DOI: 10.48420/14877285.v3 (2022). Available at: https://figshare.manchester.ac.uk/articles/dataset/Fifty_hyperspectral_reflectance_images_of_outdoor_scenes/14877285.

27. Efron, B. Better bootstrap confidence intervals. J. Am. statistical Assoc. 82, 171–185 (1987).

