## Supplementary Information for "Theoretical physical color gamuts define luminosity and naturalness perceptual limits in natural scenes"

### Supplementary figure

This section presents additional information regarding the target of Experiment 2. The shading applied to the target to create the impression of 3D sphere is illustrated in Supplementary Fig. S1. The shading effect is applied to the original circle by scaling each pixel's luminance based on its distance $d$ in pixels from the highest luminance point at $(c_x + 5, c_y - 5)$, where $(c_x, c_y)$ is the center of the circle in pixels, using the formula:

$$\text{shaded} = \text{original} \times \left(1 - \frac{d}{r \cdot s}\right), \tag{S1}$$

where $r = 16$ is the radius in pixels and $s = 1.6$ is a scalar value representing the shadow strength. The gradient is presented in the form of relative luminance, which was then multiplied by the luminance set by the observer during the completion of the experimental task, in order to create the desired final shaded sphere at different luminance intensities based on observer's adjustments.

### Supplementary tables

This section presents additional information regarding the first questionnaire given to the observers at the end of Experiments 2 and 3, as described in the main manuscript. All of the information is presented in Supplementary Tables S1 and S2 for Experiments 2 and 3, respectively: Likert-style questions with the mean and standard deviation of the responses, as well as all open-ended comments that were collected through the questionnaire.

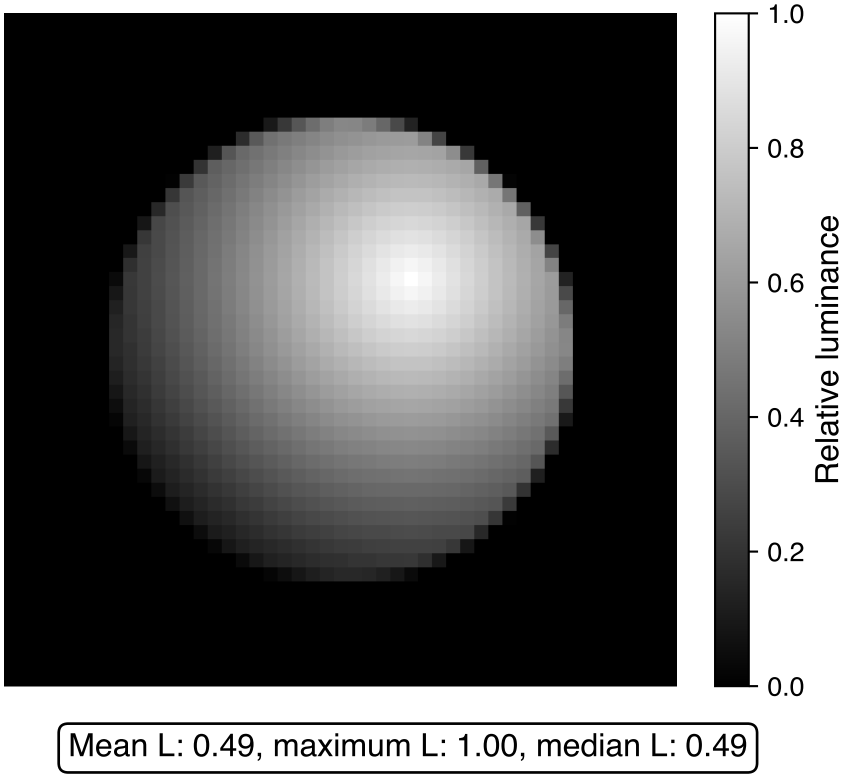


**Supplementary Figure S1.** Stimulus of Experiment 2. This represents the shading applied to the circle to create an impression of 3D sphere in Experiment 2.

| Question | Mean (± SD) |
|---|---|
| How easy was the task? 1 (difficult) to 7 (easy) | 2.75 (± 0.71) |
| Are you confident with your settings (answers)? 1 (not confident) to 7 (confident) | 4.50 (± 1.07) |
| In how many trials were you not able to find the limit of self-luminosity? 0% to 100% | 15.00 (± 14.14) |
| **Additional free comments** | |
| *Observer 1*: "The light gray and white chromaticities trials are difficult. I have to scroll up a lot and in high luminance the changes are small." | |
| *Observer 1*: "The orange-ish and blue-ish chromaticities trials are easier." | |
| *Observer 2*: "I could not find a clear threshold or limit of self-luminosity but rather a 'gray area'. I tried to find the lower limit of this gray area." | |
| *Observer 2*: "The 'gray area' between 'self-luminous' and 'illuminated surface' range is wider than in experiment 1." | |
| *Observer 3*: "There is an 'ambiguous area' between 'self-luminous' and 'illuminated surface' which is wider than in experiment 1 (where the limit was more obvious)." | |
| *Observer 3*: "The sphere does not appear as fully self-luminous but in high luminance it appears as unnatural (a portion of the area of the sphere is too bright or self-luminous and the remaining area of the sphere appears as just reflecting the light)." | |
| *Observer 4*: "It is more difficult to judge the luminosity thresholds for the 3D sphere than in experiment 1." | |
| *Observer 4*: "The white chromaticities trials are more difficult than blue and yellow." | |
| *Observer 4*: "The sphere does not seem self-luminous. I judge when the sphere seems too bright but it is difficult." | |
| *Observer 5*: "The sphere does not appear as self-luminous. It is like glossy. I judge its specular highlight as a kind of self-luminosity and tried to judge when it seemed too high." | |
| *Observer 6*: "The diffuse parts of the sphere disturb the adjustments." | |
| *Observer 6*: "The low saturation chromaticities trials are difficult." | |
| *Observer 7*: "It looks like reflecting light surface (not self-luminous). But at high luminance, it appears as unnatural. But it was difficult." | |
| *Observer 8*: "I tried to judge based on highest luminance." | |
| *Observer 8*: "I judged the point where the sphere became too bright to be in that scene." | |
| *Observer 8*: "The range is not clear and the adjustments are difficult to dose." | |

**Supplementary Table S1.** Results of first questionnaire for Experiment 2

| Question | Mean (± SD) |
|---|---|
| How easy was the task? 1 (difficult) to 7 (easy) | 5.75 (± 0.71) |
| Are you confident with your settings (answers)? 1 (not confident) to 7 (confident) | 6.25 (± 0.46) |
| In how many trials were you not able to find the limit of self-luminosity? 0% to 100% | 0.00 (± 0.00) |
| **Additional free comments** | |
| Observer 1: "It is very easy. Small adjustments are more perceptible than in experiment 2." | |
| Observer 1: "The painted wall background was difficult." | |
| Observer 2: "The target is part of the scene. It seems natural so it is easy to find the threshold. I can compare with the background. When the luminance is too high, it feels outside the image and unnatural; this is the breaking point that I judged as self-luminous even if it does not look self-luminous." | |
| Observer 3: "It is much easier than experiment 2 because the 'gray area' range is much narrower." | |
| Observer 4: "I can identify a top-down effect: I can judge from the shadows like in real life and I know that this is a natural luminance for a real object." | |
| Observer 4: "It is easy because I can separate the target from the background if the luminance is not matching. It appears as unnatural. This is the strategy that I used." | |
| Observer 5: "If the object seems self-luminous, it looks like it is outside of the picture (closer to me)." | |
| Observer 6: "The trials with the image with the small flowers were difficult." | |
| Observer 6: "The low saturation chromaticities trials were difficult." | |
| Observer 7: "The trials with the big rose were difficult." | |
| Observer 7: "Overall, it was easier because the target is inside the image." | |
| Observer 8: "I judged the self-luminosity as the limit between natural and unnatural. At high luminance, the target seems detached from the background." | |

**Supplementary Table S2.** Results of first questionnaire for Experiment 3
